# Interferon-regulated genetic programs and JAK/STAT pathway activate the intronic promoter of the short ACE2 isoform in renal proximal tubules

**DOI:** 10.1101/2021.01.15.426908

**Authors:** Jakub Jankowski, Hye Kyung Lee, Julia Wilflingseder, Lothar Hennighausen

## Abstract

Recently, a short, interferon-inducible isoform of Angiotensin-Converting Enzyme 2 (ACE2), dACE2 was identified. ACE2 is a SARS-Cov-2 receptor and changes in its renal expression have been linked to several human nephropathies. These changes were never analyzed in context of *dACE2*, as its expression was not investigated in the kidney. We used Human Primary Proximal Tubule (HPPT) cells to show genome-wide gene expression patterns after cytokine stimulation, with emphasis on the *ACE2/dACE2* locus. Putative regulatory elements controlling *dACE2* expression were identified using ChIP-seq and RNA-seq. qRT-PCR differentiating between *ACE2* and *dACE2* revealed 300- and 600-fold upregulation of *dACE2* by IFNα and IFNβ, respectively, while full length *ACE2* expression was almost unchanged. JAK inhibitor ruxolitinib ablated *STAT1* and *dACE2* expression after interferon treatment. Finally, with RNA-seq, we identified a set of genes, largely immune-related, induced by cytokine treatment. These gene expression profiles provide new insights into cytokine response of proximal tubule cells.

## INTRODUCTION

It is known that SARS-CoV-2 infectivity depends on its receptor, the Angiotensin-Converting Enzyme 2 (ACE2) (Hoffmann et al., 2020). Physiologically, ACE2 serves as an element of Renin-Angiotensin-Aldosterone system and Bradykinin system (Donoghue et al., 2000; Tipnis et al., 2000). In SARS-CoV-2 infection, the viral spike protein binds ACE2 and facilitates viral entry into cells. ACE2 expression has been detected in the kidney (Sungnak et al., 2020) and proximal tubules via single cell transcriptome analysis (Chen et al., 2020; He et al., 2020). However, transcriptional regulation of *ACE2* and its expression pattern in the kidney are poorly understood. Human studies indicate, that changes in *ACE2* expression are linked to type 2 diabetic nephropathy (Mizuiri et al., 2008), IgA nephropathy (Mizuiri et al., 2011), hypertension (Koka et al., 2008) and nephrosclerosis (Wang et al., 2010). Usually, decrease in ACE2 is associated with disease, which may dysregulate ACE/ACE2 ratio, though both ACE and ACE2 may be regulated by independent pathways (Mizuiri and Ohashi, 2015).

Acute kidney injury (AKI) is a known complication of COVID-19 and it has been proposed that decline in renal function in hospitalized patients is caused by the virus itself (Lynch and Tang, 2020). Even before the SARS-CoV-2 pandemic, AKI was a significant medical and socioeconomic burden, with estimated one in three critically ill hospitalized patients suffering from decline in kidney function (Hoste et al., 2018). If SARS-CoV-2 can indeed directly enter tubular epithelium and contribute to the injury, understanding regulation of its renal receptors, as well as associated immune response, is of utmost importance (Su et al., 2020).

Recently, a new isoform of *ACE2*, *dACE2* was identified in several cell types (Blume et al., 2020; Fignani et al., 2020; Lee et al., 2020; Ng et al., 2020; Onabajo et al., 2020). Contrary to earlier reports (Ziegler et al., 2020), where *ACE2* was suggested to be an interferon stimulated gene (ISG), expression of *dACE2* was much more significantly regulated by cytokine or viral stimulation. In fact, in come cells, like pancreatic β-cells, *dACE2* may be the prevalent isoform even at the baseline (Fignani et al., 2020). Usually, decrease in ACE2 expression is linked with disease progression, however, it is unknown whether dACE2 has an impact on these readouts, as methods used to this date assessed ACE2 without discerning between isoforms. Additionally, increased ACE2 levels were found in several animal models of kidney disease, and contribution of dACE2 to these changes remains to be assessed (Moon et al., 2008).

Here, for the first time, we investigated presence and regulation of *dACE2* in human primary proximal tubule cells. We show that *dACE2*, but not full-length *ACE2* (*flACE2*) is abundantly expressed after interferon treatment, is transcribed from an intronic promoter and contains one additional protein-coding exon. We also confirm, that similarly to lung epithelium, *dACE2* is regulated by the JAK/STAT pathway. Finally, we present a global analysis of ISGs present in renal epithelium.

These findings provide new understanding of interferon-mediated immune response in the kidney, especially in context of ACE2 activation observed in SARS-CoV-2 infection and may serve as a basis for better understanding of the commonalities between various kidney diseases.

## RESULTS AND DISCUSSION

Recent research (Blume et al., 2020; Ng et al., 2020; Onabajo et al., 2020) revealed the presence of an alternative promoter within intron 9 of the *ACE2* gene, driving expression of a short isoform of ACE2 (dACE2). Based on lung epithelium cell data, it was proposed that its extracellular enzymatic and viral spike protein-binding domains are truncated, resulting in at least partial loss of its carboxypeptidase function. While some experiments suggested the presence of *dACE2* RNA in healthy kidney tissue and tumors (Ng et al., 2020; Onabajo et al., 2020), its structure, function and presence of regulatory elements, as well as interferon-inducibility in the kidney have not been investigated in detail.

To understand the regulation of the *ACE2* locus, including *dACE2*, and to identify putative genetic control elements in human primary proximal tubules, we conducted ChIP-seq and RNA-seq experiments (Figure 1A-F) before and after IFNβ stimulation. The presence of H3K27ac (active chromatin), H3K4me1 (enhancers), H3K4me3 (promoter marks), RNA polymerase II loading (Pol II) and DNase hypersensitive sites (DHS) were used to locate putative regulatory regions.

**Figure 1.**
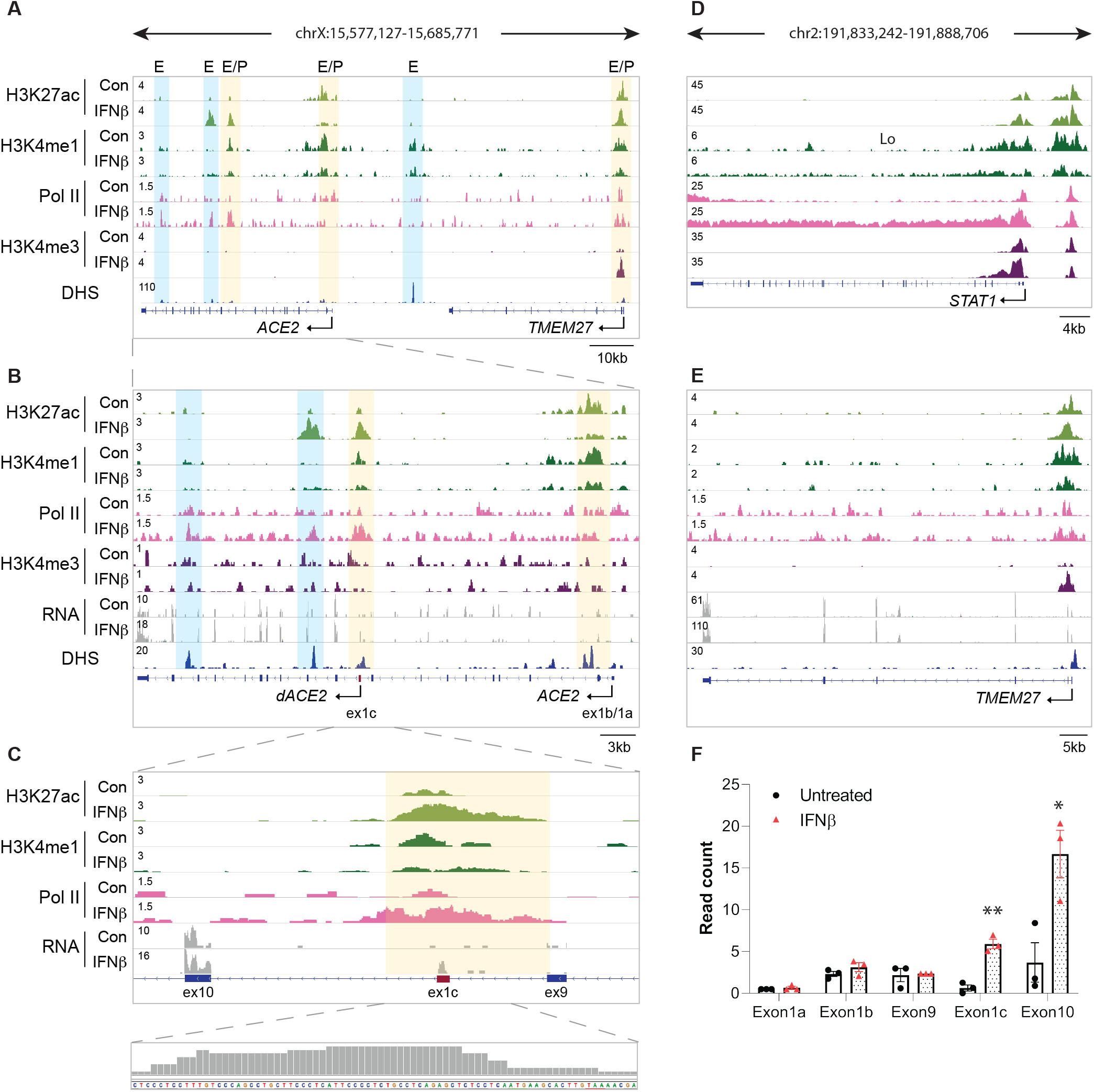
Activation of the novel intronic *ACE2* promoter by IFNβ. **(A-E)** ChIP-seq data for Pol II and histone markers H3K27ac, H3K4me1 and H3K4me3, DNase Hypersensitivity Sites and RNA-seq reads at the *ACE2, TMEM27* and *STAT1* locus in Human Primary Proximal Tubules (HTTP) with and without IFNβ. Solid arrows indicate the orientation of genes. The orange and blue shades indicate regulatory elements. **(F)** Number of RNA-seq reads at the exons 1a and 1b, and the new exon 1c. HTTP were grown in the absence or the presence of IFNβ. Individual data points as well as mean ± SEM of independent biological replicates (*n* = 3) are shown. T-test was used to assess statistical significance. \**P* < 0.05, \*\**P* < 0.01

Several such elements were identified in the *ACE2* locus (Figure 1A-C). Sequences at the recently identified promoter in intron 9 and the alternative exon 1c, the first coding exon of *dACE2*, were enriched for H3K27ac and H3K4me1 marks supporting the combined presence of promoter and enhancer elements. Interferon β (IFNβ) exposure resulted in increased H3K27ac coverage and Pol II loading, as well as increased RNA-seq read mapping demonstrating the activation of the intronic promoter (Figure 1C, F, Supplementary Figure 1). In contrast, full length *ACE2* promoter marks, which seem to be more pronounced in the kidney than in the lung(Lee et al., 2020), were decreased after IFNβ stimulation. To confirm the presence of *dACE2*, we amplified and sequenced the novel *dACE2* transcript and confirmed that exon 1c is spliced to exon 10 of *ACE2* (*Ng et al., 2020; Onabajo et al., 2020*) (Supplementary Figure 1A). Two TATA-box like sequences were identified, suggesting the presence of more than one TSS associated with the intronic promoter (Supplementary Figure 1B). Additionally, strong H3K27ac marks were induced by IFNβ around exon 11 of *ACE2*. These marks are almost absent in lung cells as reported by Lee *et al.* in lung epithelium (Lee et al., 2020). In turn, two putative enhancer elements reported at the site corresponding to 3’ end of *ACE2* gene in lung cells seem to be much weaker in the kidney. RNA-seq analyses demonstrated 5-fold IFNβ-induced expression of exon 1c, compared to exon 1a that harbors the first methionine of the full length ACE2 (Figure 1F, Supplementary Figure 1C). Finally, in addition to the *ACE2* and *TMEM27* promoters, a candidate enhancer element can be seen between the two genes, as indicated by a H3K4me1 mark. Additionally, an analysis of the extended *ACE2* locus revealed that *ACE2* and *TMEM27* are under similar interferon regulation and are bordered by CTCF chromatin boundaries suggesting that they are part of a regulatory unit (Supplementary Figure 2). The TMEM27 locus displayed increased H3K27ac and H3K4me3 promoter marks indicating gene activation after IFNβ treatment (Figure 1E). This finding may have additional significance for the kidney, as *TMEM27* gene encodes collectrin, ACE2 homologue, primarily expressed in renal proximal tubule and collecting duct(Mount, 2007). Collectrin, similarly to ACE2, regulates blood pressure(Chu and Le, 2014) and amino-acid transport (Malakauskas et al., 2007). Finally, the *STAT1* locus after interferon stimulation shows increased H3K4me3 promoter marks and Polymerase II binding, also indicating gene activation (Figure 1D).

**Figure 2.**
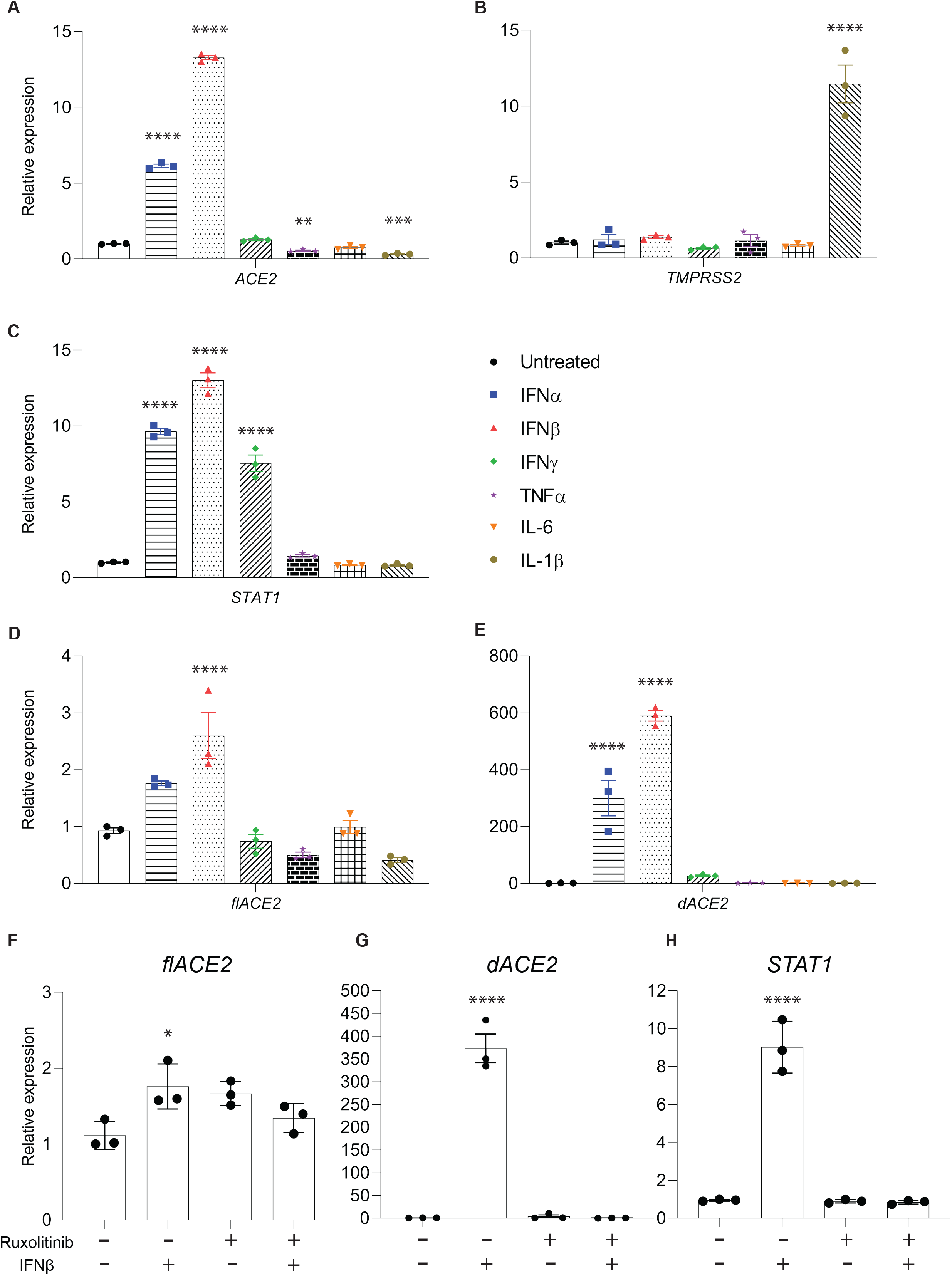
Differences in induction of total ACE2, full length ACE2 and dACE2 after cytokine treatment. **(A)** *ACE2*, *TMPRSS2* and *STAT1* mRNA levels from control and experimental cells were measured by qRT–PCR and normalized to *GAPDH* levels. Relative mRNA levels of **(D)** full length *ACE2* (*flACE2*), **(E)** *dACE2*, after cytokine treatment. **(F)** *flACE2*, **(G)** *dACE2* and **(H)** *STAT1* in cells treated with JAK inhibitor ruxolitinib or vehicle, alone or together with IFNß. Individual data points as well as mean ± SEM of independent biological replicates (*n* = 3) are shown. One- or two-way ANOVA followed by Tukey’s multiple comparisons test was used to evaluate the statistical significance of differences relative to untreated cells. \**P* < 0.05, \*\*\*\**P* < 0.0001

Next, we used qRT-PCR to assess quantitative differences between *ACE2* and *dACE2* upregulation after cytokine treatment. First, we analyzed the expression of total *ACE2* (*flACE2* and *dACE2* combined), the serine protease *TMPRSS2* for S protein priming and transcription factor *STAT1* for JAK/STAT pathway activation (Figure 2 A-C). Total ACE2 mRNA was upregulated 6- and 13-fold by IFNα and IFNβ, respectively, while *TMPRSS2* was elevated after IL-1β treatment, indicating its regulation by an independent pathway. Both IL-1β and TMPRSS2 were reported to be downregulated in nasal basal epithelium after azithromycin treatment(Renteria et al., 2020), suggesting a potential pathway shared with proximal tubules. Further, *STAT1* expression was strongly upregulated after interferon treatment. To differentiate between expression changes of *flACE2* and *dACE*2, we used isoform-specific qRT-PCR (Figure 2 D, E). While a 3-fold elevated mRNA level was detected in qRT-PCR designed for exon 9 of *ACE2* (*flACE2*), amplification of exon 1c yielded mean 300-, 590- and 27-times upregulation of *dACE2* for IFNα, IFNβ and IFNγ treatments, respectively.

To investigate whether upregulation of ACE2 or its isoform are regulated by the JAK/STAT pathway, as suggested by upregulation of *STAT1* after interferon treatment, we used a JAK1/2 inhibitor, ruxolitinib (Figure 2 F-H). Cells were incubated with 10 μM ruxolitinib or vehicle (DMSO) with or without 10 ng/ml IFNβ. Upregulation of both d*ACE2* and *STAT1* was ablated by ruxolitinib treatment, while no significant changes to full length *ACE2* expression were observed.

We also conducted unbiased RNA-seq analyses from cytokine (IFNα, IFNβ, IFNγ and IL-1β) treated cells to investigate genetic programs and ISGs in human primary proximal tubules. 735 genes were significantly induced by IFNα, 1218 by IFNβ, 1106 by IFNγ and 2142 by IL-1β (Supplementary Tables 1-4, respectively). Immune response genes were significantly enriched in the gene set significantly upregulated by IFNβ (Supplementary Table 2, Supplementary Figure 3). Expression of key immune signature genes, including IFN-response genes, was significantly induced, especially by interferon type I treatment (Figure 3 A-H). We identified interleukins (*IL4l1, IL15*), toll-like receptors (*TLR2*, *TLR4*), Interferon Regulatory Factors (*IRF1, IRF7*), Interferon Induced Proteins (*IFIT1*, *IFIT2*, *IFIT3*, *IFI44*) and chemoattractants (*CXCL10*, *CXCL11*) as significantly upregulated by IFNα and/or IFNβ. Some of the significantly upregulated genes are known to be regulators of renal injury, but belong to divergent pathways, either driving inflammation like IRF1 or TLR4 (Wang et al., 2009; Wu et al., 2007) or attenuating it like IL4 and IL15 signaling (Eini et al., 2010; Zhang et al., 2017). Recently, Ichimura et al. (Ichimura et al., 2020) reported that KIM-1 (*HAVCR1*) might serve as an additional SARS-CoV-2 receptor. KIM-1 is usually upregulated in kidney injury models, but interferon treatment alone did not yield increase in its expression.

**Figure 3.**
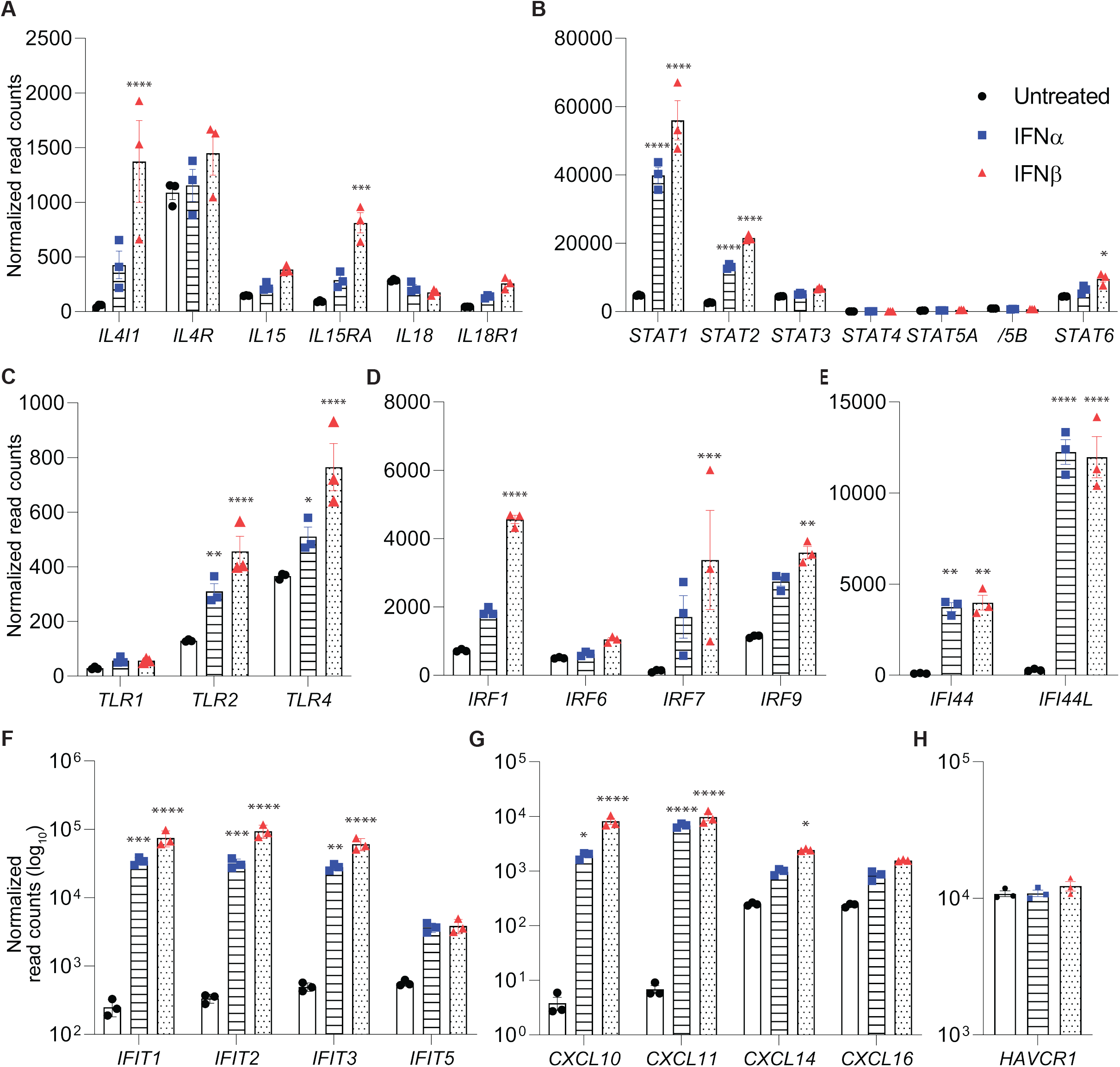
IFNα/β-induced immune response genes. **(A-H)** Relative mRNA expression levels of multiple immune genes measured by RNA-seq. Individual data points as well as mean ± SEM of independent biological replicates (*n* = 3) are shown. Significance was analyzed with one-way ANOVA followed by Tukey’s multiple comparisons test. \**P* < 0.05, \*\**P* < 0.01, \*\*\**P* < 0.001, \*\*\*\**P* < 0.0001.

While it is certain that ACE2 is necessary for SARS-Cov-2 to enter cells, our study suggests that reports of *ACE2* expression changes in response to interferon treatment and viral infection must be analyzed carefully and with *dACE2* in mind. So far, very few studies reported on *dACE2* after SARS-CoV-2 infection *in vitro*. Its role remains unknown and its promoter was postulated to be a remnant retroviral ISG (Ng et al., 2020). Blume et al.(Blume et al., 2020) report lack of increase of ACE2 or *dACE2* after SARS-CoV-2 stimulated BCi-NS1.1 lung cells. Onabajo et al. (Onabajo et al., 2020) similarly show lack of their upregulation in lung Calu3 cell line, but colon cancer Caco-2 and T84 lines exhibited slightly increased *dACE2* expression after SARS-CoV-2 exposure. This may in part be due to tissue-specific regulation of *ACE2* and *dACE2*. Standardized and validated detection method of both ACE2 isoforms, as well as understanding of regulatory elements present in *ACE2* locus is necessary to forward this topic. This is especially true for studies at the protein level, as detection methods such as Western blot are contradictory between reports (Blume et al., 2020; Ng et al., 2020). In our attempts to investigate protein levels of dACE2 using Western blot, we were able to observe a 50 kDa band, however its presence and intensity was not consistent between various anti-ACE2 antibodies (data not shown). We summarized current knowledge on factors causing dACE2 upregulation in Supplementary Table 5.

In our study, we identified several putative regulatory elements controlling *ACE2*, as well as confirmed presence of dACE2 in renal epithelium. We describe reliance of *dACE2* expression on JAK/STAT pathway, which may be of clinical importance, as JAK inhibitors are currently used to treat COVID-19(Cao et al., 2020). Finally, we present a dataset of cytokine-regulated genes in human primary proximal tubules, establishing a reference point for further studies. In conclusion, the data presented here are of relevance not only for COVID-19 pathophysiology research and therapy development, but also for diseases where ACE2 expression is altered. Despite ACE2 being a known entry gate for multiple viral strains, a clear picture of its regulation and presence in the tissues is still missing. Revisiting old paradigms and using new techniques such as ChIP- and RNA-seq may deliver novel results, as we still learn about transcriptional regulation of well-established mechanisms.

## ACKNOWLEDGMENTS

We thank Ilhan Akan, Sijung Yun and Harold Smith from the NIDDK genomics core for NGS. Cytokines were gifted by Dr. Marc Ferrer and Olive Jung (NIH/NCATS). This work utilized the computational resources of the NIH HPC Biowulf cluster (http://hpc.nih.gov).

## AUTHOR CONTRIBUTIONS

J.J., H.K.L., J.W., L.H. planned the study. J.J. carried out the experiments. H.K.L. performed computational analysis. J.J., H.K.L. analyzed the data. J.J., H.K.L., J.W., L.H. interpreted the data. J.J, H.K.L prepared the figures., J.J., H.K.L., L.H. prepared the manuscript. All authors approved final version of the manuscript.

## METHODS

### Cell culture and cytokine stimulation

Human Primary Proximal Tubule (HPPT) Cells (ATCC^®^ PCS-400-010^™^) were cultured in low-serum medium consisting of Renal Epithelial Cell Basal Medium (ATCC^®^ PCS-400-030™) with Renal Epithelial Cell Growth Kit (ATCC^®^ PCS-400-040™), Penicillin-Streptomycin-Amphotericin B Solution (ATCC^®^ PCS-999-002™) and Phenol Red (ATCC^®^ PCS-999-001™) added. Cells were obtained at passage 2, cultured according to manufacturer’s instructions and used between passages 4 and 6. In addition to characteristic cobblestone growth pattern when confluent, cells were confirmed to express several proximal tubule markers including y-glutamyltransferase-1 and HAVCR1 (KIM1) as assessed with RNA-seq. Cells were stimulated with IFNα, IFNβ, IFNγ, TNFα, IL-6 and IL-1β (all obtained from Peprotech) for 12 hours in concentration of 10 ng/ml. Cells were treated with ruxolitinib (Peprotech) at 10 μM for 12 hours together with IFNβ. At least three biological replicates were prepared for all experiments.

### RNA isolation and quantitative real-time PCR

After cytokine stimulation, cells were washed twice with PBS before RNA isolation to remove medium and debris. mRNA was isolated using PureLink™ RNA Mini Kit (Invitrogen) and 500 ng was transcribed into cDNA using SuperScript™ III First-Strand Synthesis SuperMix (Invitrogen). qRT-PCR reaction was prepared with SsoAdvanced Universal Probes Supermix (Bio-Rad) and following Taqman probes (ThermoFisher): *GAPDH* (Hs02786624_g1), *ACE2* (Hs01085333_m1), *TMPRSS2* (Hs01122322_m1) and *STAT1* (Hs01013996_m1) (Bio-Rad) or following primers: *dACE2* forward: 5’ GGAAGCAGGCTGGGACAAA 3’, *dACE2* reverse: 5’ AGCTGTCAGGAAGTCGTCCATT 3’, ACE2 forward: 5’ GGGCGACTTCAGGATCCTTAT 3’, ACE2 reverse: 5’ GGATATGCCCCATCTCATGATG 3’. Custom qRT-PCR probe sequences were as follows: ACE2: 5’ [6~FAM] ATGGACGACTTCCTGACAG [MGBE~Q] 3’, *dACE2*: 5’ [6~FAM] AGGGAGGATCCTTATGTG [MGBE~Q] 3’. Reaction conditions were as follows: initial denaturation for 3 minutes at 95°C and 40 cycles of 10 seconds at 95°C and 30 seconds at 60°C.

### PCR amplification

ACE2 PCR was performed with cDNA obtained as described above. 50ng of cDNA was used in the following reaction: initial denaturation – 3 minutes, 98°C and 35 cycles of denaturation – 30 seconds at 98°C, annealing – 30 seconds at 58°C, extension – 72°C for 2 minutes, ending with final extension of 72°C for 10 minutes. Amplified fragments were run on a 1.5% agarose in 1xTAE gel with 100 kb DNA ladder to assess product size. Bands were cut out and PCR products cleaned with MinElute Gel Extraction Kit (Quiagen) and Sanger sequenced by Quintara Biosciences. Primers used: dACE2 forward: 5’-TGTGAGAGCCTTAGGTTGGATTCC-3’, dACE2 reverse: 5’-TCTCTCCTTGGCCATGTTGT-3’.(Onabajo et al., 2020)

### RNA-seq library preparation and data analysis

mRNA was prepared as described above and quality assessed with Bioanalyzer 2100 (Aligent). Samples with adequate RIN values were transcribed into libraries using TruSeq total RNA Library Prep Kit according to manufacturer’s instructions. Libraries were pooled in equimolar amounts and sequenced with HiSeq 2000 (Illumina).

The raw data were subjected to QC analyses using the FastQC tool (version 0.11.9) (https://www.bioinformatics.babraham.ac.uk/projects/fastqc/). Trimmomatic (version 0.36) (Bolger et al., 2014) was used to assess total RNA-seq read quality and STAR RNA-seq (version 2.5.4a) (Dobin et al., 2013) using 50bp paired-end mode was used to align the reads (hg19). HTSeq (Anders et al., 2015) was used to retrieve the raw counts and R package DESeq2 (Huber et al., 2015; Love et al., 2014) was used to normalize data. Additionally, the RUVSeq (Risso et al., 2014) package was applied to remove confounding factors. Minimum of ten reads was an additional basis for filtering artifacts. The visualization was done using dplyr (https://CRAN.R-project.org/package=dplyr) and ggplot2 (Risso et al., 2014). Significantly differential expressed genes with an adjusted p-value (pAdj, FDR) below 0.05 and a fold change > 2 for up-regulated genes were categorized using GSEA (https://www.gsea-msigdb.org/gsea/msigdb). Sequence read numbers were calculated using Samtools (Li et al., 2009) software with sorted bam files.

### ChIP-seq library preparation and data analysis

Cells were washed twice with PBS and fixed with 0.75% formaldehyde in DMEM for 10 minutes in room temperature. Next, glycine was added to quench fixation in a final concentration of 125 mM and plates were incubated in room temperature for another 10 minutes. Cells were then scraped and centrifuged at 4°C, 1 minute, 3000 rpm, then washed twice with cold PBS. Pellets were re-suspended in 2 ml Farnham Lysis Buffer with protease inhibitors and incubated on ice for 10 minutes. Then, the cells were pelleted again at 4°C, 5 minutes, 3500 rpm, and re-suspended in TE buffer with protease inhibitors. Chromatin was sonicated for 3 minutes with a probe sonicator (Active Motif). Finally, after centrifugation at 4°C, 13000 g for 10 minutes, supernatant was used for immunoprecipitation.

Briefly, 600-1000 μg chromatin was incubated with antibody-coated Dynabeads™ Protein A (Invitrogen) at 4°C overnight. The beads were then washed with RIPA, high-NaCl RIPA, LiCl buffer and PBS. Next, DNA was eluted from the beads and reverse-crosslinked by incubating with proteinase K at 65°C overnight. DNA was then purified with MinElute PCR Purification Kit (Quiagen) and library preparation was performed according to manufacturer’s instructions for NEBNext^®^ Ultra™ II DNA Library Prep Kit for Illumina^®^ (New England Biotechnology). Proper library size distribution, with peak in 300-500bp range was confirmed using Bioanalyzer 2100 (Aligent), libraries pooled and sequenced with HiSeq 2000 (Illumina). Antibodies used: anti-Trimethyl-Histone H3 (Lys4) (Millipore, CS200580), Anti-RNA polymerase II CTD repeat (Abcam, ab5408), Anti-Histone H3K27ac (Active Motif, 39133), Anti-Histone H3K4me1 (Active Motif, 39297)

Quality filtering and alignment of the raw reads was done using Trimmomatic(Bolger et al., 2014) (version 0.36) and Bowtie(Langmead et al., 2009) (version 1.1.2), with the parameter ‘-m 1’ to keep only uniquely mapped reads, using the reference genome hg19. Picard tools (Broad Institute. Picard, http://broadinstitute.github.io/picard/. 2016) was used to remove duplicates and subsequently, and Homer(Heinz et al., 2010) (version 4.8.2) software was applied to generate bedGraph files. Integrative Genomics Viewer(Thorvaldsdottir et al., 2013) was used for visualization.

### Statistics

Statistical analysis of data was performed with Prism 8. First, normal distribution of data was assessed. Next, statistical significance was evaluated with 1-way or 2-way AVOVA followed by Tukey’s multiple comparisons or a T-test depending on experimental setup. Values of: \**p* < 0.05, \*\**p* < 0.01, \*\*\**p* < 0.001, \*\*\*\**p* < 0.0001 were considered statistically significant.

### Data availability

DHS from Human Renal Cortical Epithelial cells and CTCF ChIP-seq from HEK293 cells were obtained under GSE29692 and GSE68976, respectively. Hi-C data from human adrenal gland tissues was obtained from Hi-C data browser (http://3dgenome.fsm.northwestern.edu/view.php). The RNA-seq and ChIP-seq data from human primary proximal tubule cells were submitted to GEO under GSE161917 (ChIP-seq - GSE161915, RNA-seq - GSE161916), with the access token ‘yjcpuiimtlellgr’.

## SUPPLEMENTARY MATERIAL

**Supplementary Table 1.** List of all genes with normalized read counts in each replicate at Control and IFNα treated Human Primary Proximal Tubules (HTTP), log_2_ (fold change), *p*-value and adjusted *p*-value as well as upregulated gene list and GSEA analysis.

**Supplementary Table 2.** List of all genes with normalized read counts in each replicate at Control and IFNβ treated Human Primary Proximal Tubules (HTTP), log_2_ (fold change), *p*-value and adjusted *p*-value as well as upregulated gene list and GSEA analysis.

**Supplementary Table 3.** List of all genes with normalized read counts in each replicate at Control and IFNγ treated Human Primary Proximal Tubules (HTTP), log_2_ (fold change), *p*-value and adjusted *p*-value as well as upregulated gene list and GSEA analysis.

**Supplementary Table 4.** List of all genes with normalized read counts in each replicate at Control and IL-1β treated Human Primary Proximal Tubules (HTTP), log_2_ (fold change), *p*-value and adjusted *p*-value as well as upregulated gene list and GSEA analysis.

**Supplementary Table 5.** List of studies on dACE2 upregulation in multiple cell lines.

**Supplementary Figure 1.** A novel short transcript of *ACE2* was induced by interferons. **(A)** Sanger sequencing showed sequence at exon/exon boundaries of short ACE2 transcript exon1c-10. Amino acid translation is shown below. **(B)** The sequence of the first exon of dACE2. TATA boxes and TSS marked in blue and red, respectively, and CDS showed in the highlighted yellow. **(C)** IGV plot displayed RNA reads covering the entire *ACE2* gene from control and experimental cells.

**Supplementary Figure 2.** Sub-TADs defined by Hi-C from human adrenal gland and ChIP-seq data from HEK293 cells in the 470kb locus including ACE2 and neighboring genes. mRNA levels of genes in the locus under the control of IFNβ measured by RNA-seq.

**Supplementary Figure 3.** Percentage of induced gene classes via Hallmark Gene Sets (FDR q-value < 0.05).

## REFERENCES

Anders, S., Pyl, P.T., and Huber, W. (2015). HTSeq--a Python framework to work with high-throughput sequencing data. Bioinformatics 31, 166–169.

Blume, C., Jackson, C.L., Spalluto, C.M., Legebeke, J., Nazlamova, L., Conforti, F., Perotin-Collard, J.-M., Frank, M., Crispin, M., Coles, J., et al. (2020). A novel isoform of ACE2 is expressed in human nasal and bronchial respiratory epithelia and is upregulated in response to RNA respiratory virus infection. bioRxiv, 2020.2007.2031.230870.

Bolger, A.M., Lohse, M., and Usadel, B. (2014). Trimmomatic: a flexible trimmer for Illumina sequence data. Bioinformatics 30, 2114–2120.

Cao, Y., Wei, J., Zou, L., Jiang, T., Wang, G., Chen, L., Huang, L., Meng, F., Huang, L., Wang, N., et al. (2020). Ruxolitinib in treatment of severe coronavirus disease 2019 (COVID-19): A multicenter, single-blind, randomized controlled trial. J Allergy Clin Immunol 146, 137–146 e133.

Chen, Q.L., Li, J.Q., Xiang, Z.D., Lang, Y., Guo, G.J., and Liu, Z.H. (2020). Localization of Cell Receptor-Related Genes of SARS-CoV-2 in the Kidney through Single-Cell Transcriptome Analysis. Kidney diseases (Basel, Switzerland) 6, 258–270.

Chu, P.L., and Le, T.H. (2014). Role of collectrin, an ACE2 homologue, in blood pressure homeostasis. Curr Hypertens Rep 16, 490.

Dobin, A., Davis, C.A., Schlesinger, F., Drenkow, J., Zaleski, C., Jha, S., Batut, P., Chaisson, M., and Gingeras, T.R. (2013). STAR: ultrafast universal RNA-seq aligner. Bioinformatics 29, 15–21.

Donoghue, M., Hsieh, F., Baronas, E., Godbout, K., Gosselin, M., Stagliano, N., Donovan, M., Woolf, B., Robison, K., Jeyaseelan, R., et al. (2000). A novel angiotensin-converting enzyme-related carboxypeptidase (ACE2) converts angiotensin I to angiotensin 1-9. Circ Res 87, E1–9.

Eini, H., Tejman-Yarden, N., Lewis, E.C., Chaimovitz, C., Zlotnik, M., and Douvdevani, A. (2010). Association between renal injury and reduced interleukin-15 and interleukin-15 receptor levels in acute kidney injury. J Interferon Cytokine Res 30, 1–8.

Fignani, D., Licata, G., Brusco, N., Nigi, L., Grieco, G.E., Marselli, L., Overbergh, L., Gysemans, C., Colli, M.L., Marchetti, P., et al. (2020). SARS-CoV-2 receptor Angiotensin I-Converting Enzyme type 2 (ACE2) is expressed in human pancreatic β-cells and in the human pancreas microvasculature. bioRxiv, 2020.2007.2023.208041.

He, Q., Mok, T.N., Yun, L., He, C., Li, J., and Pan, J. (2020). Single-cell RNA sequencing analysis of human kidney reveals the presence of ACE2 receptor: A potential pathway of COVID-19 infection. Molecular genetics & genomic medicine, e1442.

Heinz, S., Benner, C., Spann, N., Bertolino, E., Lin, Y.C., Laslo, P., Cheng, J.X., Murre, C., Singh, H., and Glass, C.K. (2010). Simple combinations of lineage-determining transcription factors prime cis-regulatory elements required for macrophage and B cell identities. Mol Cell 38, 576–589.

Hoffmann, M., Kleine-Weber, H., Schroeder, S., Kruger, N., Herrler, T., Erichsen, S., Schiergens, T.S., Herrler, G., Wu, N.H., Nitsche, A., et al. (2020). SARS-CoV-2 Cell Entry Depends on ACE2 and TMPRSS2 and Is Blocked by a Clinically Proven Protease Inhibitor. Cell 181, 271–280 e278.

Hoste, E.A.J., Kellum, J.A., Selby, N.M., Zarbock, A., Palevsky, P.M., Bagshaw, S.M., Goldstein, S.L., Cerda, J., and Chawla, L.S. (2018). Global epidemiology and outcomes of acute kidney injury. Nat Rev Nephrol 14, 607–625.

Huber, W., Carey, V.J., Gentleman, R., Anders, S., Carlson, M., Carvalho, B.S., Bravo, H.C., Davis, S., Gatto, L., Girke, T., et al. (2015). Orchestrating high-throughput genomic analysis with Bioconductor. Nat Methods 12, 115–121.

Ichimura, T., Mori, Y., Aschauer, P., Padmanabha Das, K.M., Padera, R.F., Weins, A., Nasr, M.L., and Bonventre, J.V. (2020). KIM-1/TIM-1 is a Receptor for SARS-CoV-2 in Lung and Kidney. medRxiv.

Koka, V., Huang, X.R., Chung, A.C., Wang, W., Truong, L.D., and Lan, H.Y. (2008). Angiotensin II up-regulates angiotensin I-converting enzyme (ACE), but down-regulates ACE2 via the AT1-ERK/p38 MAP kinase pathway. Am J Pathol 172, 1174–1183.

Langmead, B., Trapnell, C., Pop, M., and Salzberg, S.L. (2009). Ultrafast and memory-efficient alignment of short DNA sequences to the human genome. Genome Biol 10, R25.

Lee, H.K., Jung, O., and Hennighausen, L. (2020). Regulation of the *ACE2* locus in human airways cells. bioRxiv, 2020.2010.2004.325415.

Li, H., Handsaker, B., Wysoker, A., Fennell, T., Ruan, J., Homer, N., Marth, G., Abecasis, G., Durbin, R., and Genome Project Data Processing, S. (2009). The Sequence Alignment/Map format and SAMtools. Bioinformatics 25, 2078–2079.

Love, M.I., Huber, W., and Anders, S. (2014). Moderated estimation of fold change and dispersion for RNA-seq data with DESeq2. Genome Biol 15, 550.

Lynch, M.R., and Tang, J. (2020). COVID-19 and Kidney Injury. Rhode Island medical journal (2013) 103, 24–28.

Malakauskas, S.M., Quan, H., Fields, T.A., McCall, S.J., Yu, M.J., Kourany, W.M., Frey, C.W., and Le, T.H. (2007). Aminoaciduria and altered renal expression of luminal amino acid transporters in mice lacking novel gene collectrin. Am J Physiol Renal Physiol 292, F533–544.

Mizuiri, S., Hemmi, H., Arita, M., Aoki, T., Ohashi, Y., Miyagi, M., Sakai, K., Shibuya, K., Hase, H., and Aikawa, A. (2011). Increased ACE and decreased ACE2 expression in kidneys from patients with IgA nephropathy. Nephron Clin Pract 117, c57–66.

Mizuiri, S., Hemmi, H., Arita, M., Ohashi, Y., Tanaka, Y., Miyagi, M., Sakai, K., Ishikawa, Y., Shibuya, K., Hase, H., et al. (2008). Expression of ACE and ACE2 in individuals with diabetic kidney disease and healthy controls. Am J Kidney Dis 51, 613–623.

Mizuiri, S., and Ohashi, Y. (2015). ACE and ACE2 in kidney disease. World J Nephrol 4, 74–82.

Moon, J.Y., Jeong, K.H., Lee, S.H., Lee, T.W., Ihm, C.G., and Lim, S.J. (2008). Renal ACE and ACE2 expression in early diabetic rats. Nephron Exp Nephrol 110, e8–e16.

Mount, D.B. (2007). Collectrin and the kidney. Curr Opin Nephrol Hypertens 16, 427–429.

Ng, K.W., Attig, J., Bolland, W., Young, G.R., Major, J., Wrobel, A.G., Gamblin, S., Wack, A., and Kassiotis, G. (2020). Tissue-specific and interferon-inducible expression of nonfunctional ACE2 through endogenous retroelement co-option. Nat Genet.

Onabajo, O.O., Banday, A.R., Stanifer, M.L., Yan, W., Obajemu, A., Santer, D.M., Florez-Vargas, O., Piontkivska, H., Vargas, J.M., Ring, T.J., et al. (2020). Interferons and viruses induce a novel truncated ACE2 isoform and not the full-length SARS-CoV-2 receptor. Nat Genet.

Renteria, A.E., Endam Mfuna, L., Adam, D., Filali-Mouhim, A., Maniakas, A., Rousseau, S., Brochiero, E., Gallo, S., and Desrosiers, M. (2020). Azithromycin Downregulates Gene Expression of IL-1beta and Pathways Involving TMPRSS2 and TMPRSS11D Required by SARS-CoV-2. Am J Respir Cell Mol Biol.

Risso, D., Ngai, J., Speed, T.P., and Dudoit, S. (2014). Normalization of RNA-seq data using factor analysis of control genes or samples. Nat Biotechnol 32, 896–902.

Su, H., Yang, M., Wan, C., Yi, L.X., Tang, F., Zhu, H.Y., Yi, F., Yang, H.C., Fogo, A.B., Nie, X., et al. (2020). Renal histopathological analysis of 26 postmortem findings of patients with COVID-19 in China. Kidney Int 98, 219–227.

Sungnak, W., Huang, N., Becavin, C., Berg, M., Queen, R., Litvinukova, M., Talavera-Lopez, C., Maatz, H., Reichart, D., Sampaziotis, F., et al. (2020). SARS-CoV-2 entry factors are highly expressed in nasal epithelial cells together with innate immune genes. Nat Med 26, 681–687.

Thorvaldsdottir, H., Robinson, J.T., and Mesirov, J.P. (2013). Integrative Genomics Viewer (IGV): high-performance genomics data visualization and exploration. Brief Bioinform 14, 178–192.

Tipnis, S.R., Hooper, N.M., Hyde, R., Karran, E., Christie, G., and Turner, A.J. (2000). A human homolog of angiotensin-converting enzyme. Cloning and functional expression as a captopril-insensitive carboxypeptidase. J Biol Chem 275, 33238–33243.

Wang, G., Kwan, B.C., Lai, F.M., Choi, P.C., Chow, K.M., Li, P.K., and Szeto, C.C. (2010). Intrarenal expression of miRNAs in patients with hypertensive nephrosclerosis. Am J Hypertens 23, 78–84.

Wang, Y., John, R., Chen, J., Richardson, J.A., Shelton, J.M., Bennett, M., Zhou, X.J., Nagami, G.T., Zhang, Y., Wu, Q.Q., et al. (2009). IRF-1 promotes inflammation early after ischemic acute kidney injury. J Am Soc Nephrol 20, 1544–1555.

Wu, H., Chen, G., Wyburn, K.R., Yin, J., Bertolino, P., Eris, J.M., Alexander, S.I., Sharland, A.F., and Chadban, S.J. (2007). TLR4 activation mediates kidney ischemia/reperfusion injury. J Clin Invest 117, 2847–2859.

Zhang, M.Z., Wang, X., Wang, Y., Niu, A., Wang, S., Zou, C., and Harris, R.C. (2017). IL-4/IL-13-mediated polarization of renal macrophages/dendritic cells to an M2a phenotype is essential for recovery from acute kidney injury. Kidney Int 91, 375–386.

Ziegler, C.G.K., Allon, S.J., Nyquist, S.K., Mbano, I.M., Miao, V.N., Tzouanas, C.N., Cao, Y., Yousif, A.S., Bals, J., Hauser, B.M., et al. (2020). SARS-CoV-2 Receptor ACE2 Is an Interferon-Stimulated Gene in Human Airway Epithelial Cells and Is Detected in Specific Cell Subsets across Tissues. Cell 181, 1016–1035 e1019.

