## Supplementary Figure 1 for "Interferon-regulated genetic programs and JAK/STAT pathway activate the intronic promoter of the short ACE2 isoform in renal proximal tubules"

A

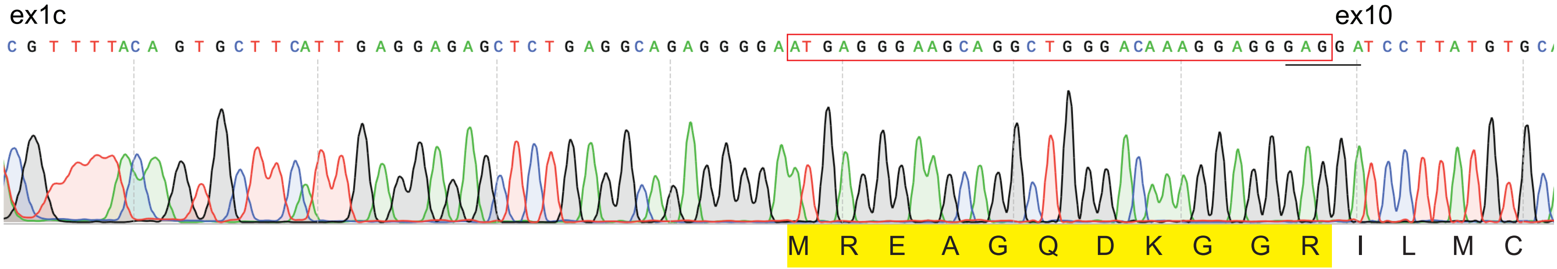


B

new promoter and exon1c

GACAAGGTA**TTTAA**CTTTCTTTGGTTTCAGTTTCCTTATTT**TATAAA**GTAGAATAGTAATTCCCAGGTTGCAGGCTTGTGAGA
GCCTTAGGTTGGATTCCCTAGCTTGAAAAGGAGATCGTTTTACAAGTGCTTCATTGAGGAGAGCTCTGAGGCAGAGGGGA
ATGAGGGAAGCAGGCTGGGACAAAGGAGGGAG

C

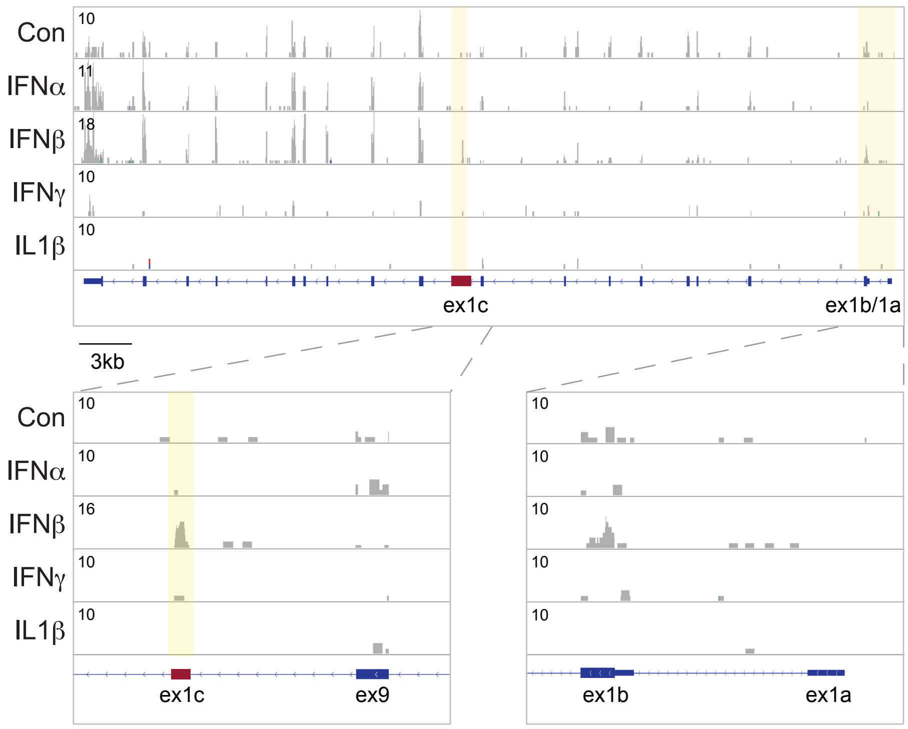
