## Supplementary figures and images for "Interferon-regulated genetic programs and JAK/STAT pathway activate the intronic promoter of the short ACE2 isoform in renal proximal tubules"

### Supplementary Figure 2

A

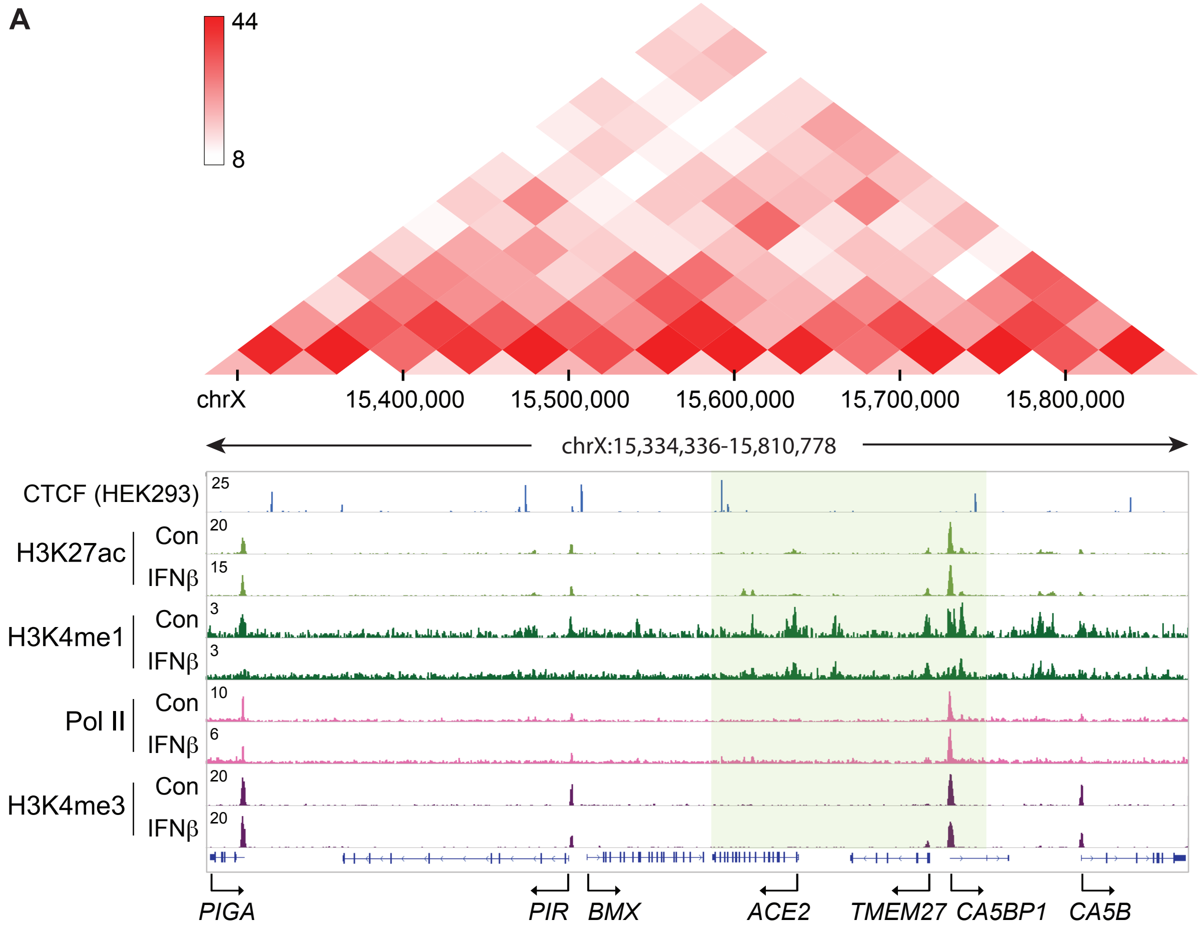


| RNA-seq | *PIGA* | *PIR* | *BMX* | *ACE2* | *TMEM27* | *CA5BP1* | *CA5B* |
|---|---|---|---|---|---|---|---|
| Con | 372.6 | 24.9 | 0.9 | 20.2 | 114.5 | 130.2 | 51.0 |
| IFNβ | 388.9 | 21.6 | 2.3 | 155.6 | 499.0 | 105.9 | 57.8 |
| Induction | 1.0 | 0.9 | 2.6 | 7.7 | 4.4 | 0.8 | 1.1 |

### Supplementary Figure 3

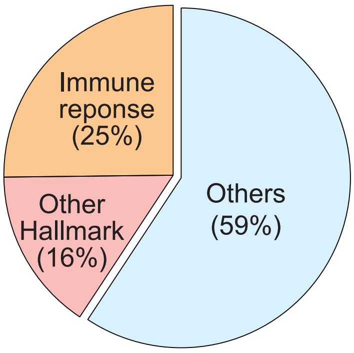
